# Ciliopathic micrognathia is caused by aberrant skeletal differentiation and remodeling

**DOI:** 10.1101/2020.06.19.162073

**Authors:** Christian Louis Bonatto Paese, Evan C. Brooks, Megan Aarnio-Peterson, Samantha A. Brugmann

## Abstract

Ciliopathies represent a growing class of diseases caused by defects in microtubule-based organelles called primary cilia. Approximately 30% of ciliopathies can be characterized by craniofacial phenotypes such as craniosynostosis, cleft lip/palate and micrognathia. Patients with ciliopathic micrognathia experience a particular set of difficulties including impaired feeding and breathing and have extremely limited treatment options. To understand the cellular and molecular basis for ciliopathic micrognathia, we utilized the *talpid*^*2*^ (*ta^2^*), a bona fide avian model for the human ciliopathy Oral-Facial-Digital syndrome subtype 14 (OFD14). Histological analyses revealed that the onset of ciliopathic micrognathia in *ta*^*2*^ embryos occurred at the earliest stages of mandibular development. Neural crest-derived skeletal progenitor cells were particularly sensitive to a ciliopathic insult, undergoing unchecked passage through the cell cycle and subsequent increased proliferation. Furthermore, whereas neural crest-derived skeletal differentiation was initiated, osteoblast maturation failed to progress to completion. Additional molecular analyses revealed that an imbalance in the ratio of bone deposition and resorption also contributed to ciliopathic micrognathia in *ta*^*2*^ embryos. Thus, our results suggest that ciliopathic micrognathia is a consequence of multiple, aberrant cellular processes necessary for skeletal development, and provide potential avenues for future therapeutic treatments.

## Introduction

Primary cilia are ubiquitous microtubule-based organelles that serve as signaling hubs for multiple molecular signaling pathways (Goetz and Anderson, 2010). Disruptions in the structure or function of primary cilia result in a class of disorders known as ciliopathies. Clinically, ciliopathies commonly present with a range of phenotypes including polydactyly, situs inversus, retinitis pigmentosa, and renal cystic disease (Baker and Beales, 2009). Approximately 30% of all ciliopathies can be classified as craniofacial ciliopathies due to the craniofacial complex being the primary organ system affected (Zaghloul and Brugmann, 2011). Craniofacial ciliopathies are characterized by the common presentation of several phenotypes including cleft palate, craniosynostosis, and micrognathia (Schock and Brugmann, 2017). While some progress has been made into the study of craniofacial ciliopathies (Cela et al., 2018; Chang et al., 2016a; Chang et al., 2014; Kawasaki et al., 2017; Millington et al., 2017; Schock et al., 2015; Tian et al., 2017; Watanabe et al., 2019), the cellular and molecular etiologies of ciliopathic craniofacial skeletal anomalies remains unclear.

Development of the mandible begins with the generation of cranial neural crest cells (NCCs) from the rostral neural tube. A subset of Hox-negative NCCs migrate from the dorsal neural tube into the first branchial arch where they proliferate and populate the mandibular prominence (MNP) (Couly et al., 1996; Kontges and Lumsden, 1996). After initial formation and patterning of the MNP, NCCs begin to differentiate into skeletal derivatives. A subset of NCCs condense and differentiate into chondrocytes to form a bilateral cartilaginous structure called Meckel’s cartilage. Meckel’s cartilage serves as a template for proper growth of the mandible, but it is not necessary for mandibular bone development (Mori-Akiyama et al., 2003). Other NCCs in the MNP undergo intramembranous ossification, in which NCCs condense into compact nodules that differentiate directly into osteoblasts. During intramembranous ossification, osteoblast differentiation is initiated when *Distal-less homeobox 5* (*DLX5*) induces the expression of *Runt related transcription factor 2* (*RUNX2*), the master transcriptional regulator of bone development (Holleville et al., 2007; Lee et al., 2003). *RUNX2*+ NCCs differentiate into osteoblasts and secrete a specialized extracellular matrix called osteoid tissue. Osteoblasts embedded within the osteoid mature into osteocytes (reviewed in (Franz-Odendaal, 2011).

After bone deposition, the developing mandible begins to take its characteristic shape via bone resorption. Within the craniofacial skeleton, both mesoderm-derived osteoclasts and neural crest-derived osteocytes contribute to bone resorption via the secretion of tartrate-resistant acid phosphatase (TRAP) (Minkin, 1982; Qing et al., 2012; Tang et al., 2012), osteoclast-specific *Matrix metalloproteinase 9* (*MMP9*) (Engsig et al., 2000; Reponen et al., 1994), or osteocyte-specific *Matrix metalloproteinase 13* (*MMP13*) (Behonick et al., 2007; Johansson et al., 1997). TRAP is secreted into the bony matrix where it dephosphorylates the structural phosphoproteins osteopontin and bone sialoprotein (Ek-Rylander et al., 1994), while matrix metalloproteinases (MMPs) enzymatically degrade extracellular matrix components such as collagen and elastin to remodel tissues (reviewed in (Cui et al., 2017)). Levels of bone resorption are inversely proportional to jaw size and play an important role in determining overall mandibular size between and among species (Ealba et al., 2015). Despite numerous studies demonstrating that ciliary dysfunction leads to micrognathia (Adel Al-Lami et al., 2016; Brugmann et al., 2010; Cela et al., 2018; Gray et al., 2009; Kitamura et al., 2020; Kolpakova-Hart et al., 2007; Zhang et al., 2011), if or how loss of ciliary function affects bone resorption has yet been described.

Due to ample *in ovo* accessibility of NCCs and conserved organization of facial prominences, avian embryos have long been utilized to study craniofacial development (reviewed in (Schock et al., 2016)). Improved genome sequencing coupled with the existence of naturally occurring avian mutants have recently allowed researchers to make significant advances in understanding the etiology of developmental disorders using an avian model system. Perhaps the most utilized mutant avian lines have been those of the *talpid* family (Abbott et al., 1959; Abbott et al., 1960; Ede and Kelly, 1964a; Ede and Kelly, 1964b). *talpid*^*2*^ (*ta^2^*) is a naturally occurring avian mutant that is characterized by its striking presentation of polydactyly and craniofacial phenotypes (Abbott et al., 1959; Abbott et al., 1960; Brugmann et al., 2010; Chang et al., 2014; Dvorak and Fallon, 1992; Munoz-Sanjuan et al., 2001; Schneider et al., 1999). Our previous studies revealed that the causative mutation in the *ta*^*2*^ was a 19 bp deletion in *C2 calcium-dependent domain containing 3* (*C2CD3*) (Chang et al., 2014), a distal centriolar protein coding gene required for ciliogenesis (Hoover et al., 2008; Ye et al., 2014). Impaired C2CD3-dependent ciliogenesis in the *ta*^*2*^ results in facial clefting, ectopic archosaurian-like first generation teeth, hypo- or aglossia, and micrognathia (Chang et al., 2014; Chang et al., 2016b; Harris et al., 2006; Munoz-Sanjuan et al., 2001; Schock et al., 2015). Genetic, molecular and bioinformatic data from these studies determined that *ta*^*2*^ was a bona fide model for the human craniofacial ciliopathy, Oral-facial-digital syndrome 14 (OFD14) (Schock et al., 2015). While micrognathia is a common and severe craniofacial phenotype associated with OFD14 (Boczek et al., 2018; Cortes et al., 2016; Thauvin-Robinet et al., 2014), the molecular and cellular etiology of micrognathia in OFD14 patients and in *ta*^*2*^ embryos has yet to be explored.

In this study, we present the first in-depth analysis of the onset of ciliopathic micrognathia using the *ta*^*2*^ model. Our data demonstrate that although there is an expansion of the SOX9+ osteochondroprogenitor population in the *ta^2^*, osteoprogenitors undergo precocious and incomplete differentiation, resulting in a reduced number of mature osteoblasts. Furthermore, analysis of bone remodeling markers demonstrated that there is increased bone resorption in the *ta*^*2*^ mandible. Thus, our data suggest that ciliopathic micrognathia is due to the combinatorial effect of aberrant skeletal differentiation and remodeling.

## Results

### *ta^2^* embryos present with micrognathia and dysmorphic skeletal elements

To understand the etiology of ciliopathic micrognathia, we first characterized the onset of micrognathia in *ta*^*2*^ embryos. The mandibular skeleton was first detectable via Alizarin Red staining at Hamburger-Hamilton stage 36 (HH36). At this stage, skeletal elements that comprise the mandible, including the angular, dentary, splenial and surangular bones, were visible (Fig. 1A). Relative to controls, Alizarin Red staining of HH36 *ta*^*2*^ mandibles revealed a reduction in calcified tissue (Fig. 1B). Volumetric measurements of the *ta*^*2*^ angular, dentary, and surangular bones confirmed a significant reduction in size relative to analogous control skeletal elements (Fig. 1C). Although the overall volume of the *ta*^*2*^ splenial bone was not significantly different from the control at this stage (Fig. S1A), it was reduced in length and appeared to have a medial, ectopic skeletal element via Alizarin Red (Fig. 1B) and microCT analysis (Fig. 1D-E). While total mandibular volume and surface area were not significantly different at this stage (Fig. S1B-C), total length measurements revealed that the *ta*^*2*^ mandibles were significantly shorter than control mandibles at HH36 (Fig. 1F).

**Fig. 1.**
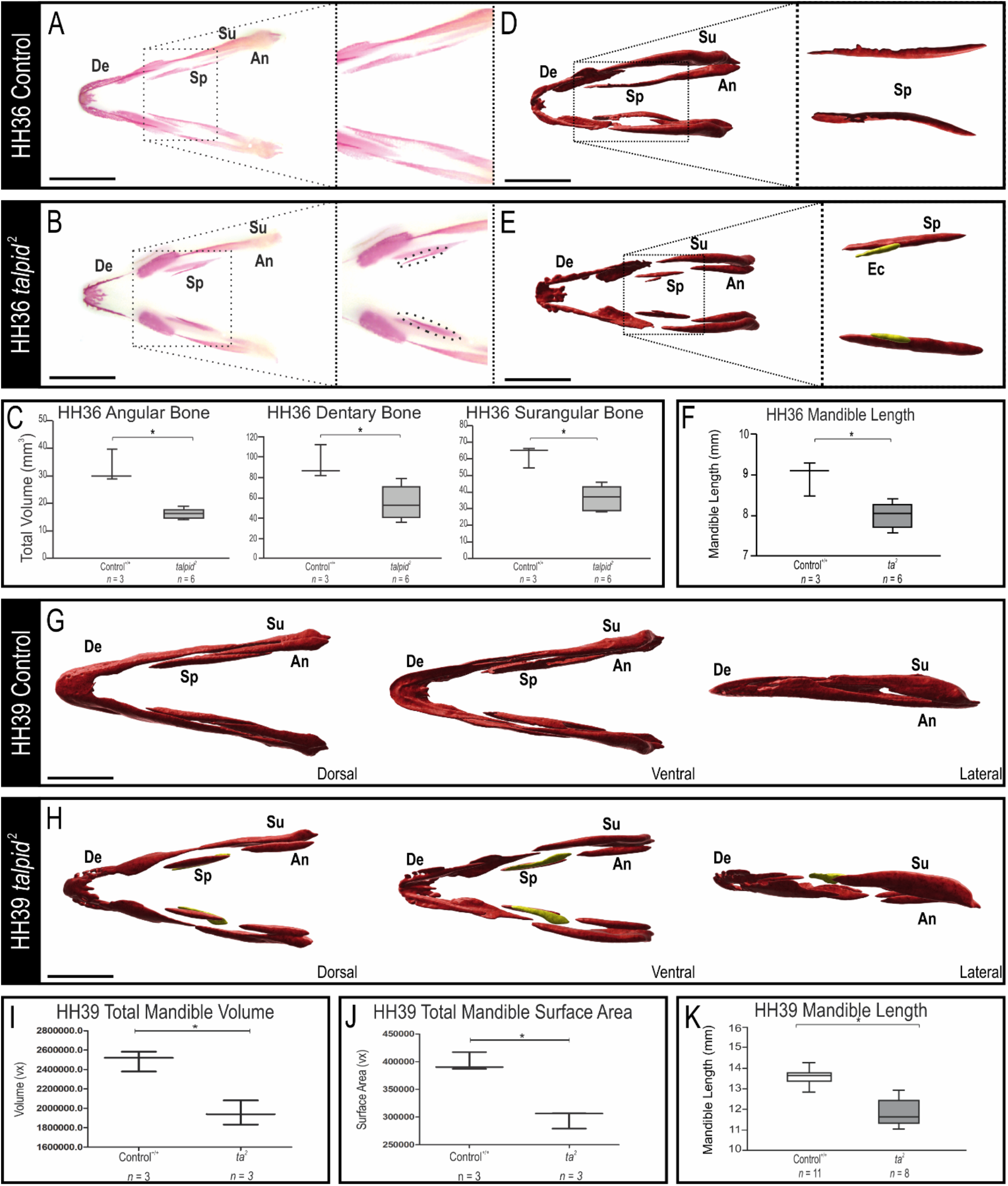
Characterization of micrognathic onset in *ta*^*2*^ embryos. (A-B) Ventral views of Alizarin Red stained HH36 control^+/+^ (n = 3) and *ta*^*2*^ (n = 6) mandibles, with insets showing a higher magnification of the splenial bones. Dotted lines in (B) inset denote duplication of the splenial bone. (C) Volumetric measurements of HH36 control^+/+^ and *ta*^*2*^ angular, dentary and surangular bones. (D-E) Ventral views of microCT scans of HH36 control^+/+^ and *ta*^*2*^ mandibles. Insets show higher magnification images of splenial bones (n = 3 per group). For better visualization, the ectopic bone is pseudocolored in yellow. (F) Length measurements of HH36 control^+/+^ and *ta*^*2*^ mandibles. (G-H) Various views of microCT scans of HH39 control^+/+^ and *ta*^*2*^ mandibles (n= 3 per group). For better visualization, the ectopic bone is pseudocolored in yellow. (I) Total volume and (J) surface area measurements of HH39 control^+/+^ and *ta*^*2*^ mandibles. (K) Length measurements of HH39 control^+/+^ and *ta*^*2*^ mandibles. An: Angular, De: Dentary, Ec: Ectopic splenial bone, Sp: Splenial, Su: Surangular, Vx: Voxels. Data are mean ± s.d. Scale bars: 1cm (A-B, D-E and G-H). Statistical analysis was performed by student’s t-test (* denotes *P*< 0.05).

We further characterized mandibular development in control and *ta*^*2*^ embryos at HH39, which is the latest stage we can consistently harvest *ta*^*2*^ embryos before embryonic lethality (Abbott et al., 1959; Abbott et al., 1960). At this stage, the developing *ta*^*2*^ mandible was reduced both in volume and surface area when compared to stage-matched control mandibles (Fig. 1G-J). microCT analysis confirmed the presence of an ectopic skeletal element medially adjacent to a severely reduced splenial bone in HH39 *ta*^*2*^ mandibles (Fig. 1H, yellow). These findings were further supported by wholemount skeletal staining with Alizarin Red and Alcian Blue (Fig. S1D-E). Compared to HH39 control embryos, stage-matched *ta*^*2*^ mandibles were dysmorphic and possessed reduced Alizarin Red staining, indicative of reduced bone deposition (Fig. S1E). Length measurements between controls and stage-matched *ta*^*2*^ mandibles confirmed that the micrognathia first observed at HH36 persisted in HH39 embryos (Fig. 1K).

In addition to analyzing the skeletal elements of the mandible proper, we also analyzed skeletal elements of the *ta*^*2*^ tongue since hypoglossia is present in both *ta*^*2*^ embryos and OFD14 patients (Boczek et al., 2018; Chang et al., 2014; Schock et al., 2015; Thauvin-Robinet et al., 2014). Avians have a distinct glossal apparatus that contains osteogenic and cartilaginous elements supported by the lingual process of the hyoid bone and rudimentary lingual muscles. Unlike mandibular bones that form through intramembranous ossification, the avian tongue and hyoid apparatus have elements that form through endochondral ossification (Lillie and Hamilton, 1952). Length measurements of whole mount Alizarin Red and Alcian Blue stained control and *ta*^*2*^ ceratobranchial bones demonstrated that the *ta*^*2*^ glossal apparatus was also reduced in size relative to controls, suggesting that both endochondral and intramembranous ossification were affected in *ta*^*2*^ mutants (Fig. S1F-H). Considering these findings, we next sought to examine the etiology of ciliopathic micrognathia by examining both cellular processes and molecular pathways required for mandibular development.

### Cell cycle progression and cell proliferation are perturbed in *ta^2^* mandibular prominences

Development of the mandibular skeleton is dependent upon expansion of NCCs within the developing MNP. Expansion of any cell population requires proliferation and proper progression through the cell cycle. Ciliopathic mutants are particularly vulnerable to cell cycle disruptions as the centrioles required for ciliogenesis are the same organelles required for formation of the mitotic spindle. Thus, ciliary extension and cell cycle progression are inextricably linked processes (Fig. 2A) (Tucker et al., 1979). To examine if cell cycle progression is impaired in *ta*^*2*^ MNPs, we examined expression of three specific markers of the G1/S checkpoint, a point at which primary cilium retraction is necessary for cell passage into S phase and DNA replication (Tucker et al., 1979). First, p21 arrests the passage from G1 to S-phase if DNA damage or microtubular aberration is detected. If an alteration is not detected, the cell cycle progress to the S-phase (Barr et al., 2017; el-Deiry et al., 1993). Second, Cyclin E binds to CDK2 to drive the transition from G1 to S phase (reviewed in (Woo and Poon, 2003). Lastly, to control for aberrant apoptosis, p21 inhibits the expression of the pro-apoptotic factor Survivin (Suzuki et al., 2000). We quantified the expression of *p21*, *cyclin E* and *survivin* in HH26 control and *ta*^*2*^ MNPs. Quantitative reverse transcription PCR (qRT-PCR) analysis revealed *p21* expression was significantly decreased (48% downregulated) in the *ta*^*2*^ MNPs relative to controls, whereas *cyclin E* and *survivin* were significantly increased (44% and 21%, respectively) (Fig. 2B). These results suggested that the loss of C2CD3-dependent ciliogenesis resulted in unrestricted cell-cycle progression within cells of the *ta*^*2*^ MNP.

**Fig. 2.**
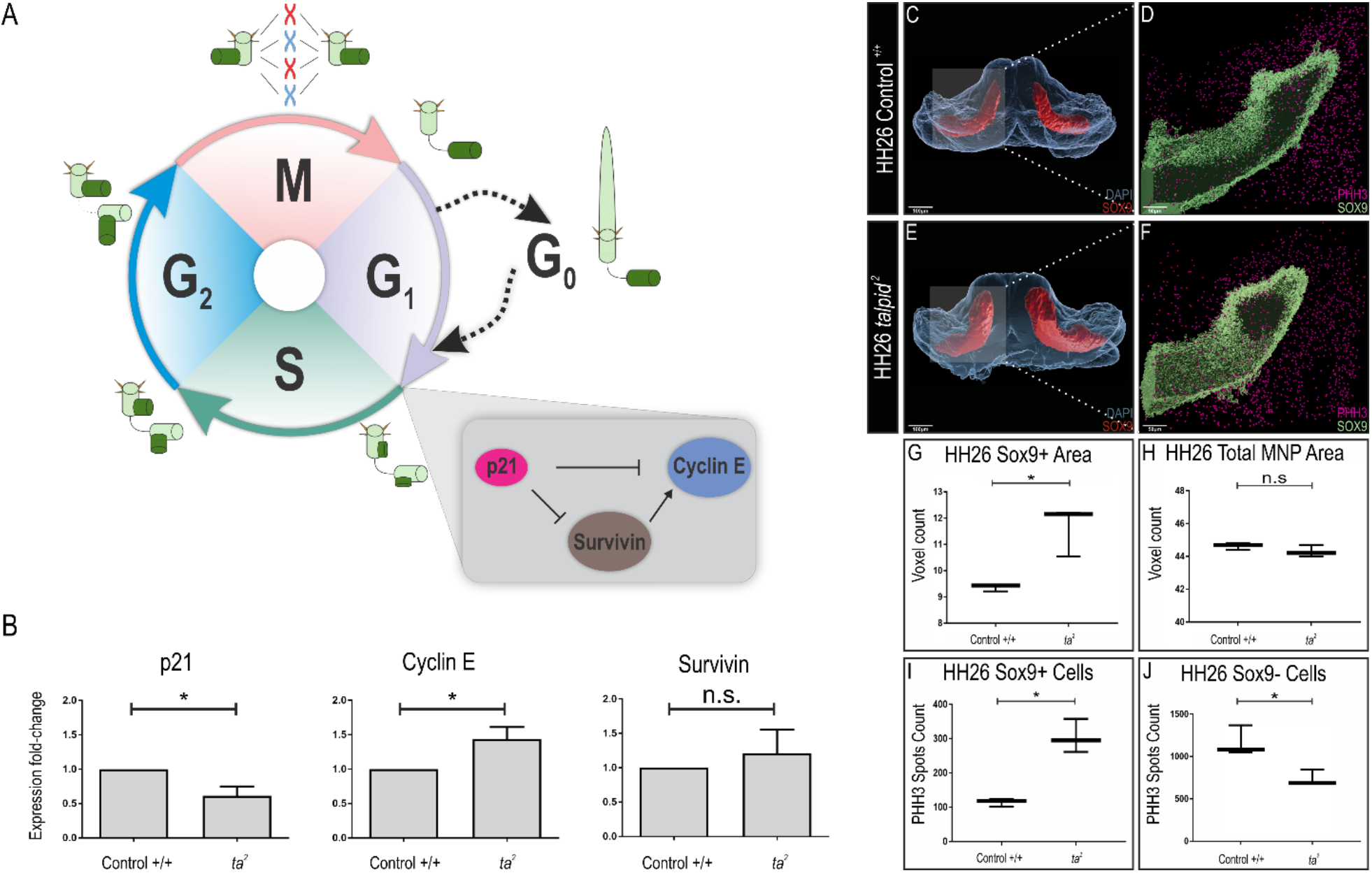
Cell cycle progression and proliferation are impaired in NCC-derived skeletal progenitors within the *ta*^*2*^ MNPs. (A) Schematic diagram of the cell cycle and primary cilia assembly. The grey inset highlights the gene regulatory network at the G1/S-phase checkpoint. (B) qRT-PCR for *p21*, *cyclin E*, and *survivin* in HH29 control^+/+^ and *ta*^*2*^ MNP (n = 3 for control^+/+^ and *ta*^*2*^ samples). (C-F) Immunostaining for SOX9 (red) and DAPI (blue) surfaces in HH26 (C) control^+/+^ and (E) *ta*^*2*^ MNPs. Immunostaining for PHH3 (purple) and SOX9 (green) in HH26 (D) control^+/+^ and (F) *ta*^*2*^ MNPs. (G) Voxel count of SOX9+ cells in HH26 control^+/+^ and *ta*^*2*^ MNPs (n = 3). (H) Voxel count of DAPI surface in MNPs of HH26 control^+/+^ and *ta*^*2*^ (n = 3). (I) PHH3+ puncta counts in the SOX9+ cell population of control^+/+^ and *ta*^*2*^ MNPs (n = 3). (J) PHH3 spots count in the SOX9-cell population of HH26 control^+/+^ and *ta*^*2*^ MNPs (n = 3). Data are mean ± s.d. Scale bars are (C, E)100μm and (D,F) 50μm. Statistical analysis was performed by student’s t-test (* denotes *P*< 0.005).

Given these results, we next sought to examine cell proliferation and apoptosis rates in the osteochondroprogenitor population of the *ta*^*2*^ MNP. Immunostaining for SOX9 (Ng et al., 1997) and phosphorylated Histone H3 (PHH3) in HH26 control and *ta*^*2*^ MNPs revealed that *ta*^*2*^ MNPs have an increased SOX9+ population relative to control MNPs (Fig. 2C-G), despite the overall area of the MNP remaining constant (Fig. 2H). Double immunostaining for PHH3 and SOX9 further revealed that proliferation was significantly increased specifically within the SOX9+ population (Fig. 2I), but not SOX9-mesenchyme (Fig. 2J). Cell death analysis revealed apoptosis was significantly decreased in *ta*^*2*^ MNPs across both SOX9+ and SOX9-populations (Fig. S2A-F), which correlates with *survivin* expression in these tissues. Thus, these data suggested that there was aberrant proliferation specifically within SOX9+ osteochondroprogenitor cells in the *ta*^*2*^ MNP.

### Increased *RUNX2* expression results in an increased preosteoblast population in *ta^2^* mandibles

After sufficient proliferation, multipotent NCCs differentiate into several lineages, including the chondrogenic and osteogenic derivatives (Le Douarin, 1982). *RUNX2* functions as the master transcriptional regulator for osteoblast differentiation, as increased *RUNX2* expression is required for the initial skeletal differentiation of NCCs into preosteoblasts (Komori et al., 1997; Otto et al., 1997). Once NCCs express *RUNX2*, they proceed down the osteoblast lineage (Fig. 3A) (Kobayashi et al., 2000). In the MNP, *RUNX2* expression is regulated both positively and negatively by additional transcription factors expressed in the developing MNP. *DLX5* induces the expression of *RUNX2* (Holleville et al., 2007; Lee et al., 2003), whereas *Heart and neural crest derivatives expressed 2* (*HAND2*) inhibits *RUNX2* expression (Funato et al., 2009). To determine if osteoblast differentiation was disrupted in *ta*^*2*^ MNPs, we spatially and quantitatively examined expression of these genes. Both whole-mount and single molecule fluorescent *in situ* hybridization (Wang et al., 2012) revealed that *DLX5* expression was expanded distally around Meckel’s cartilage in the HH29 *ta*^*2*^ MNPs (Fig 3B-E’), relative to controls. qRT-PCR analysis further supported these findings and revealed a significant, 22% increase of *DLX5* expression in the *ta*^*2*^ MNP (Fig. 3F). Conversely, *HAND2* expression was decreased, and shifted proximally within the developing *ta*^*2*^ MNP (Fig. 3G-J’). qRT-PCR confirmed a significant, 35% decrease of *HAND2* expression in the *ta*^*2*^ MNP relative to the control MNP (Fig. 3K). Lastly, examination of *RUNX2* expression demonstrated a significant and robust proximal expansion (Fig. 3L-O’). This observation was confirmed quantitatively via qRT-PCR which revealed *RUNX2* expression was increased by 45% in the *ta*^*2*^ MNP (Fig. 3P). Sagittal Z-stack projections of the MNP revealed that transcripts for all three genes were detected ventrally adjacent to the SOX9+ population (Fig. S3A-F). Together, these data suggested that excessive expression of *RUNX2* due to the increased expression of the positive regulator *DLX5* and decreased expression of the negative regulator *HAND2* resulted in ectopic production of pre-osteoblasts in the *ta*^*2*^ MNP. We next sought to determine if the maturation of pre-osteoblasts into osteoblasts was impaired in ciliopathic micrognathia.

**Fig. 3.**
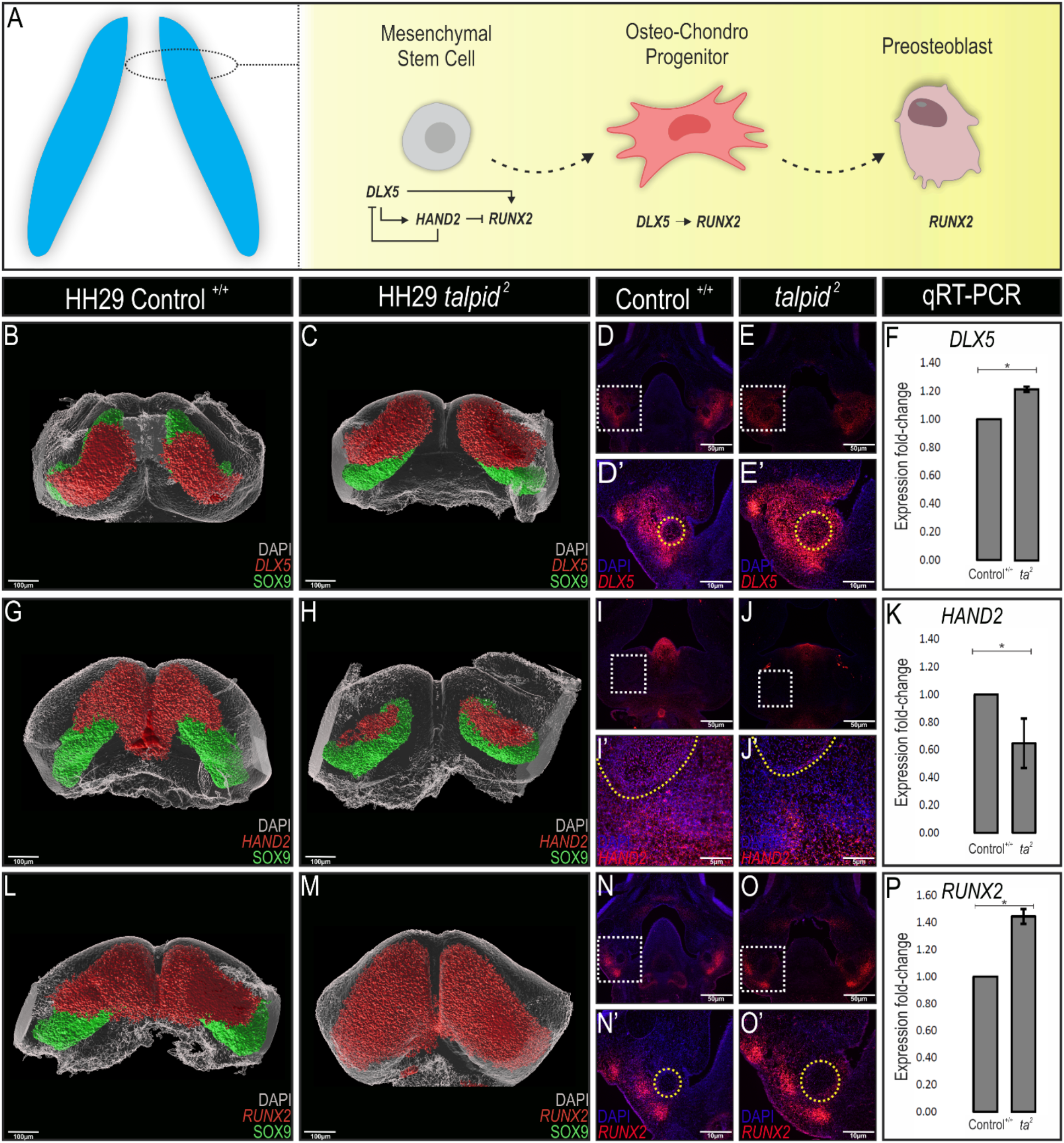
Differentiation of NCC-derived osteochondroprogenitors is impaired in *ta*^*2*^ MNPs. (A) Schematic demonstrating the osteochondroprogenitor gene regulatory network present in the developing MNP. (B-C) Wholemount *DLX5 in situ* hybridization (red) in HH29 control^+/+^ and *ta*^*2*^ MNPs counterstained with SOX9 antibody (green). (D-E’) RNAscope *in situ* hybridization for *DLX5* in frontally sectioned HH29 (D) control^+/+^ and (E) *ta*^*2*^ heads with higher magnifications in D’ and E’. (F) qRT-PCR for *DLX5* transcripts in HH29 control^+/+^ and *ta*^*2*^ MNPs (n = 3). (G-H) Wholemount *HAND2 in situ* hybridization (red) in HH29 control^+/+^ and *ta*^*2*^ MNPs counterstained with SOX9 antibody (green). (I-J’) RNAscope *in situ* hybridization for *HAND2* in frontally sectioned HH29 (I) control^+/+^ and (J) *ta*^*2*^ heads with higher magnifications in I’ and J’. (K) qRT-PCR for *HAND2* transcripts in HH29 control^+/+^ and *ta*^*2*^ MNPs (n = 3). (L-M) Wholemount *RUNX2 in situ* hybridization in HH29 control^+/+^ and *ta*^*2*^ MNPs counterstained with SOX9 antibody (green). (N-O’) RNAscope *in situ* hybridization for *RUNX2* in frontally sectioned HH29 (N) control^+/+^ and (O) *ta*^*2*^ heads with higher magnifications in N’ and O’. (P) qRT-PCR for *RUNX2* transcripts in HH29 control^+/+^ and *ta*^*2*^ MNPs (n = 3). Data are mean ± s.d. Yellow circles in D’, E’, I’, J’, N’, and O’ outline Meckel’s cartilage. Statistical analysis was performed by student’s t-test (* denotes *P*< 0.05).

### Osteoblast maturation is impaired in *ta^2^* mutants

Increased *RUNX2* expression could be indicative of increased bone deposition; however, in the *ta^2^*, increased *RUNX2* expression accompanies a micrognathic phenotype. We hypothesized that despite an expansion of pre-osteoblasts, transition towards a mature osteoblast was impaired in *ta*^*2*^ mandibles. Osteoblast maturation occurs in three stages. In the first stage of osteoblast maturation, pre-osteoblasts proliferate and express fibronectin and collagen type 1 (COL1A1) proteins. In the second stage, pre-osteoblasts exit the cell cycle and differentiate, while expressing alkaline phosphatase (ALPL) and COL1A1 (Rodrigues et al., 2012). In the last phase, mature osteoblasts express osteocalcin (OCN), a protein essential for matrix mineralization (Fig. 4A) (reviewed in (Rutkovskiy et al., 2016). To determine if osteoblast maturation was impaired, we analyzed the expression of these three markers. Immunostaining revealed increased expression of COL1A1 in the developing *ta*^*2*^ mandible when compared to controls (Fig. 4B-C). Increased expression was confirmed quantitatively via qRT-PCR (Fig. 4D). *ALPL* expression, which was localized to cortical surfaces of the developing skeletal elements in HH39 control mandibles, was mislocalized and reduced in *ta*^*2*^ mandibles (Fig. 4E-F). Quantification of *ALPL* transcripts in HH35 mandibles revealed a 35% downregulation in *ta*^*2*^ in comparison to control mandibles (Fig. 4G). Lastly, we analyzed the expression of OCN. OCN expression was reduced in the developing *ta*^*2*^ mandible when compared to controls (Fig. 4H-I). qRT-PCR analysis confirmed OCN expression was significantly reduced by approximately 72% in *ta*^*2*^ mandibles relative to controls (Fig. 4J). Taken together, these results suggest that despite an expansion of the pre-osteoblast population, osteoblast maturation was severely impaired. Thus, our data suggested that impaired osteoblast maturation and subsequently reduced bone deposition contributed to ciliopathic micrognathia.

**Fig. 4.**
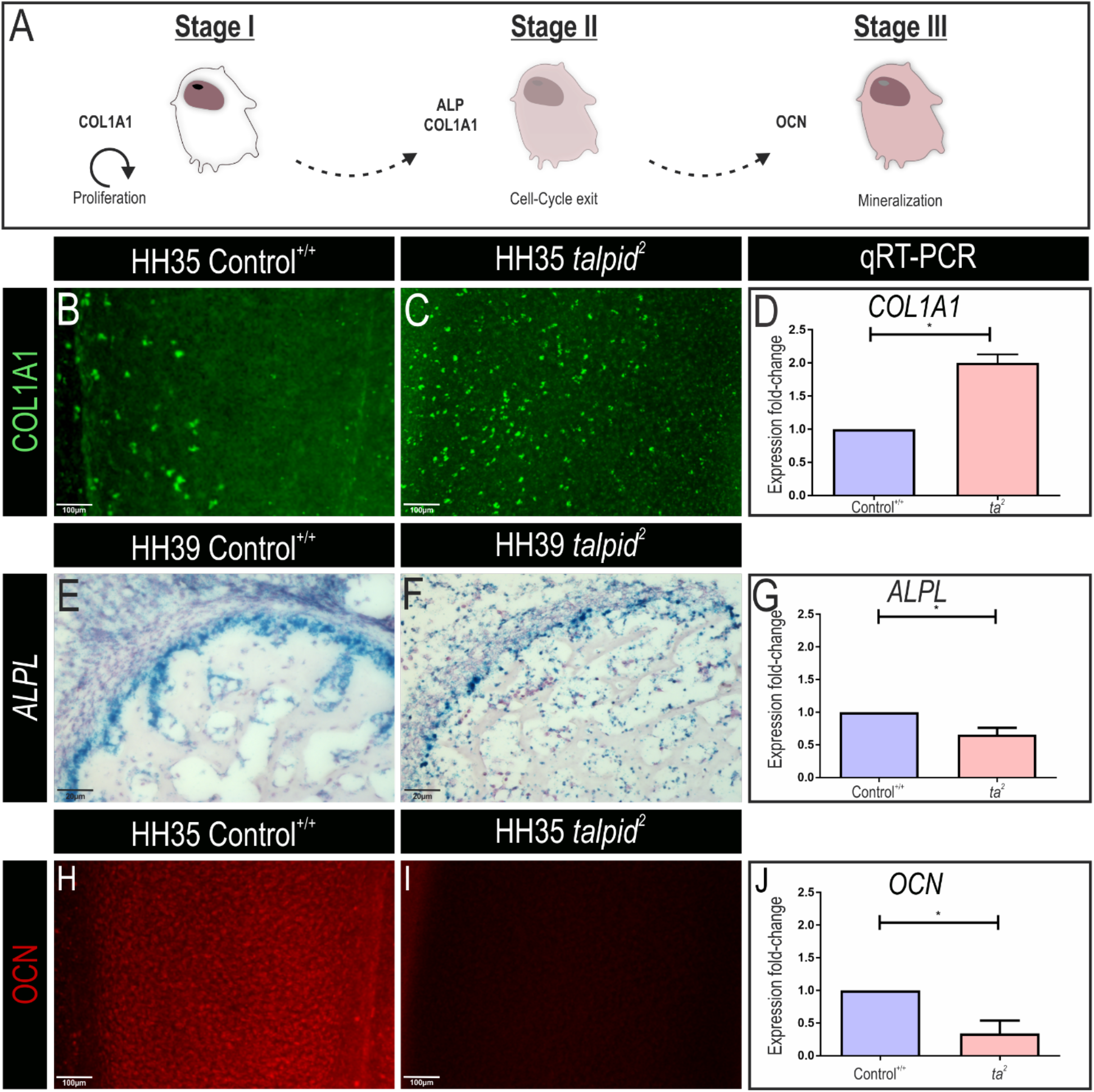
Osteoblast maturation in impaired in *ta*^*2*^ mandibles. (A) Schematic of osteoblastic maturation stages. (B-C) Wholemount immunofluorescence staining for COL1A1 in control^+/+^ and *ta*^*2*^ mandibles at HH35. (D) Quantification of *COL1A1* transcripts by qRT-PCR in HH35 control^+/+^ and *ta*^*2*^ mandibles (n = 3). (E-F) RNAscope *in situ* hybridization staining for *ALPL* in HH39 control^+/+^ and *ta*^*2*^ surangular sections denoted in blue (n= 3). (G) Quantification of *ALPL* transcripts by qRT-PCR in HH35 control^+/+^ and *ta*^*2*^ mandibles (n = 3). (H-I) Whole mount immunofluorescence staining for OCN in control^+/+^ and *ta*^*2*^ mandibles at HH35. (J) Quantification of *OCN* transcripts by qRT-PCR in HH35 control^+/+^ and *ta*^*2*^ mandibles (n = 3). Data are mean ± s.d. Statistical analysis was performed by student’s t-test (* denotes *P*< 0.05).

### Bone resorption is upregulated in *ta^2^* mandibles

Several studies have reported that proper determination of jaw length requires regulated bone remodeling. Bone remodeling is a dynamic process that consists of resorption of bony matrix by osteoclasts and osteocytes and deposition of new bony matrix by osteoblasts. We next sought to determine if aberrant bone remodeling also contributed to ciliopathic micrognathia, as it was previously reported that the amount of bone resorption was inversely proportional to jaw length in avian embryos (Ealba et al., 2015). To test the hypothesis that increased bone resorption contributed to ciliopathic micrognathia, we examined the expression of several markers for this process. Matrix metalloproteinases (MMPs) are a family of proteases necessary for osteoclast recruitment and solubilization of the osteoid during bone remodeling (reviewed in (Cui et al., 2017)). During mandibular remodeling, *MMP13* is expressed in NCC-derived osteoblasts and osteocytes (Behonick et al., 2007; Johansson et al., 1997). Both *in situ* hybridization and qRT-PCR revealed a significant increase in *MMP13* expression in the HH39 *ta*^*2*^ mandible when compared to controls (Fig. 5A-C). To confirm this result, we analyzed a second marker of bone resorption, *secreted phosphoprotein 1* (*SPP1*). *SPP1* is responsible for osteoclast adhesion and bone resorption across various species (Choi et al., 2008; Pinero et al., 1995). *SPP1* was more prominently expressed in the medullar region of the developing *ta*^*2*^ mandible relative to controls (Fig. 5D-E). These data were quantitatively verified via qRT-PCR revealing a 77% upregulation of *SPP1* expression in the *ta*^*2*^ mandibles relative to controls (Fig. 5F). Finally, we examined the ratio of osteoprotegerin (OPG) to receptor activator of NF-κB ligand (RANKL) expression. OPG protects bones from excessive resorption by binding to RANKL and preventing it from binding to its receptor, RANK (Fig. 5G). qRT-PCR analyses revealed that while OPG expression was significantly decreased, RANKL expression was significantly increased in the developing *ta*^*2*^ mandible (Fig. 5H). Aberrant expression of both markers resulted in an increased ratio of OPG/RANKL expression (Fig. 5H), further suggesting that excessive bone resorption was contributing to ciliopathic micrognathia in *ta*^*2*^ embryos.

**Fig. 5.**
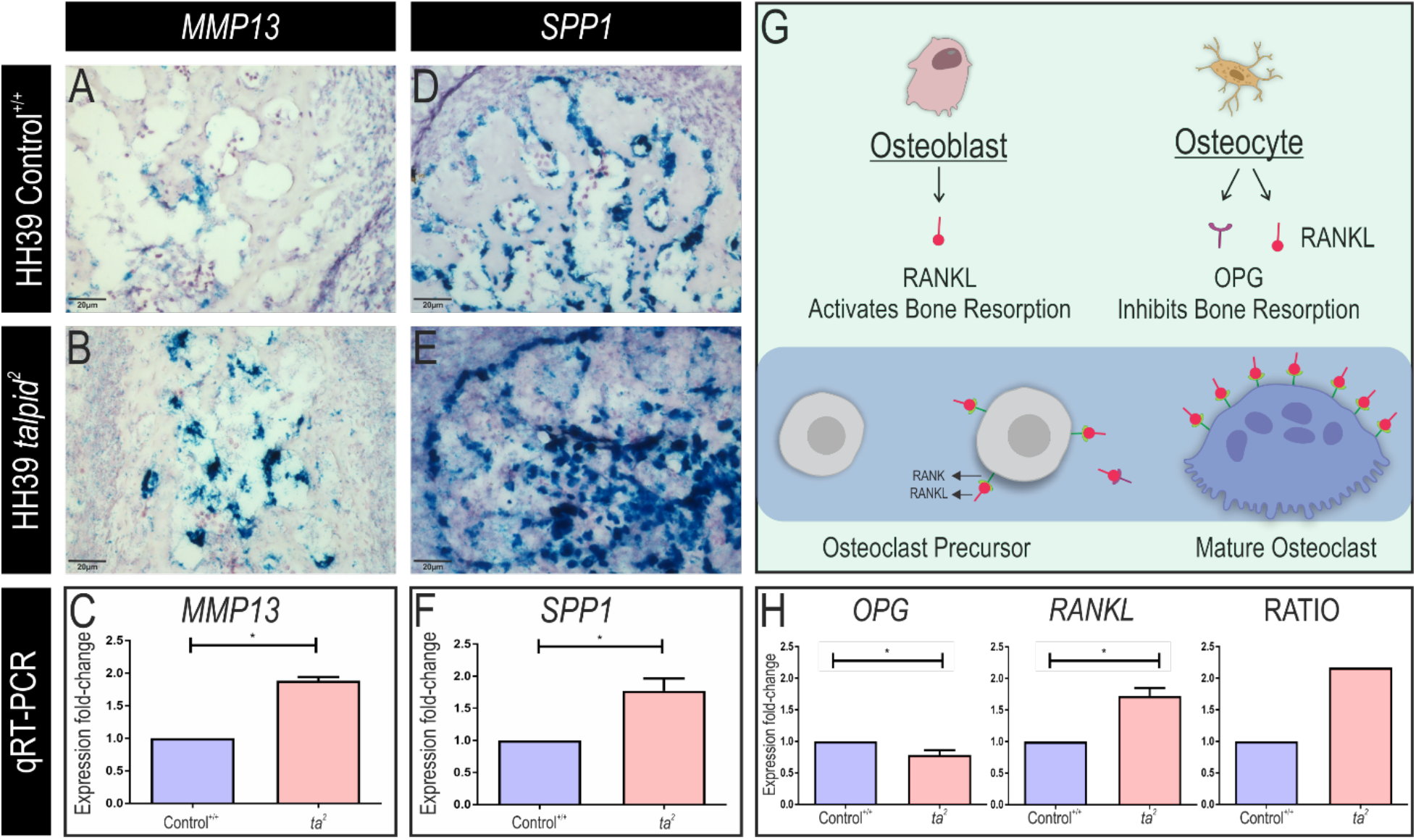
Bone resorption is increased in *ta*^*2*^ mandibles. (A-B) RNAscope *in situ* hybridization staining for *MMP13* in HH39 control^+/+^ and *ta*^*2*^ surangular sections in blue (n = 3). (C) qRT-PCR quantification of *MMP13* transcripts in HH39 control^+/+^ and *ta*^*2*^ mandibles (p < 0.05, n = 3 per group). (D-E) RNAscope *in situ* hybridization staining for *SPP1* in HH39 control^+/+^ and *ta*^*2*^ surangular sections in blue (n = 3). (F) qRT-PCR quantification of *SPP1* transcripts in HH39 control^+/+^ and *ta*^*2*^ mandibles (n = 3). (G) Schematic of RANKL/OPG regulation of osteoclast differentiation in normal development. (H) Quantification of *OPG* and *RANKL* expression in HH39 control^+/+^ and *ta*^*2*^ mandibles and its ratio (n = 3). Data are mean ± s.d. Statistical analysis was performed by student’s t-test (* denotes *P*< 0.05).

Herein, we explored the etiology of ciliopathic micrognathia using the *ta*^*2*^ model as our guide. We found that despite increased proliferation and expansion of osteochondroprogenitors and preosteoblast populations, osteoblast maturation and subsequent bone deposition was impaired in the *ta*^*2*^ mandible. Furthermore, excessive amounts of bone resorption were also detected following impaired bone deposition (Fig. 6). These studies are significant as they increase the understanding of the cellular processes and molecular pathways that can be targeted for future therapeutic approaches for ciliopathies.

**Fig. 6.**
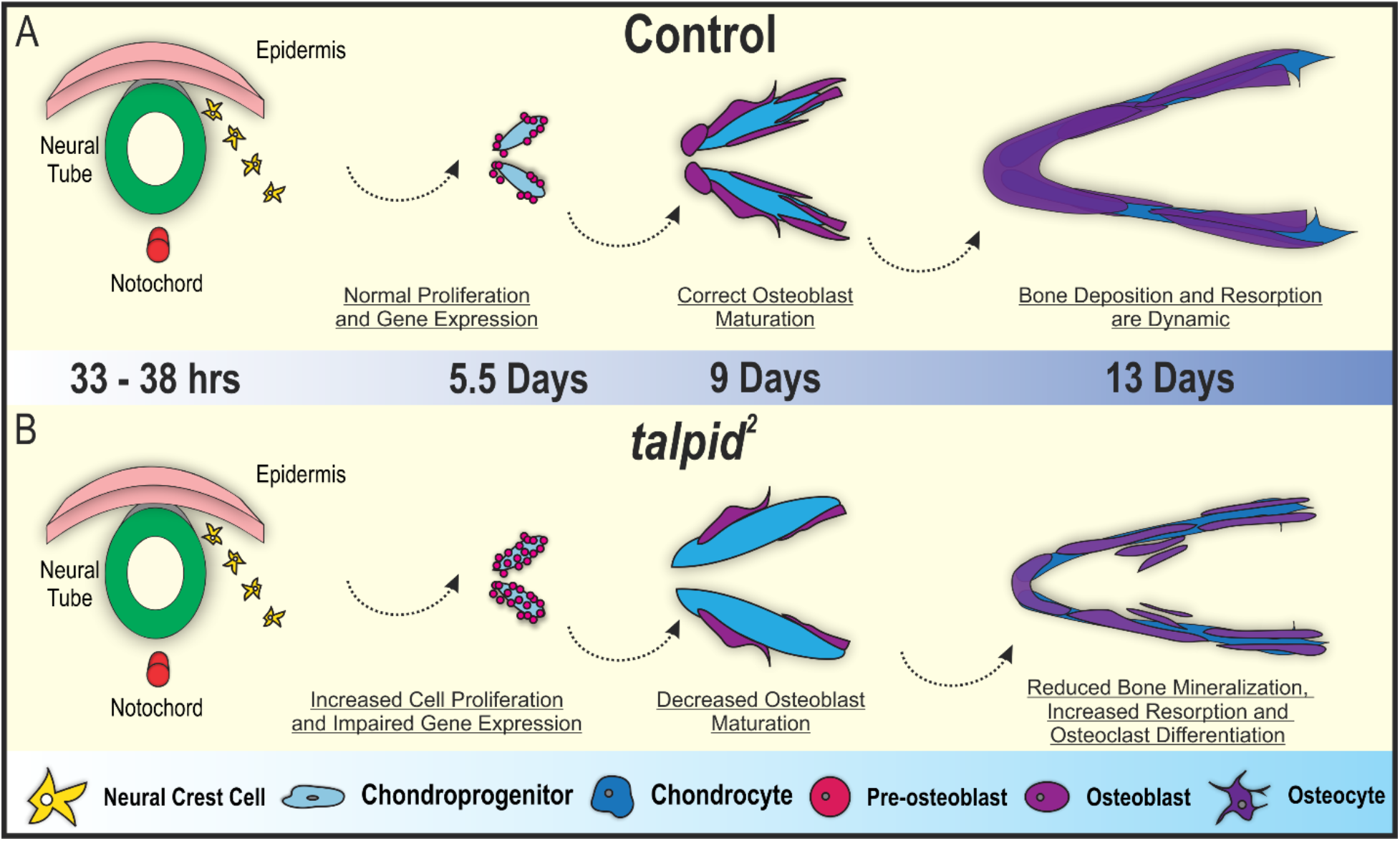
Summary of mandibular osteogenesis in control and *ta*^*2*^ embryos. (A-B) Schematic of NCC-derived intramembranous ossification that occurs in control and *ta*^*2*^ mandibles. In summary, NCCs migrate from the dorsal neural tube and populate the MNP. NCCs differentiate into osteochondroprogenitors at 5.5 days of development. In *ta*^*2*^ MNPs, impaired gene expression and increased cell proliferation lead to increased pre-osteoblast deposition in the MNPs. Markers of osteoblast maturation were dysregulated in *ta*^*2*^ MNPs at 9 days of development, causing decreased bone mineralization.

## Discussion

Micrognathia is a significant biomedical burden present in almost 20% of ciliopathies (Schock and Brugmann, 2017). The predominant treatment for micrognathia is distraction osteogenesis, an invasive and traumatic procedure in which the mandible is cut and gradually separated to allow new bone growth until the mandible reaches a desired size (Ilizarov, 1988). To develop less invasive therapeutic options for ciliopathic micrognathia, it is important to gain a more extensive understanding of its etiology. While studies have shown that loss of primary cilia leads to micrognathia, a thorough analysis of how the cellular and molecular mechanisms essential for mandibular development are impaired in ciliopathic mutants have yet to be described. To this end, we utilized the avian ciliopathic mutant *ta^2^*, a bona fide model of the human ciliopathy OFD14, to investigate how ciliopathic micrognathia arose. While the majority of ciliopathic studies utilize conditional knockout murine models, the naturally occurring *ta*^*2*^ model, in which the ciliopathic insult is ubiquitous throughout the organism, allows for analyses more relevant to human ciliopathic conditions. Our data revealed that loss of C2cd3-mediated ciliogenesis resulted in unchecked cell cycle progression, increased proliferation of SOX9+ osteochondroprogenitors and impaired expression of gene regulatory networks necessary for the maturation of NCC-derived osteoblasts. Furthermore, excessive bone resorption was detected in *ta*^*2*^ mandibles later in development (Fig. 6A-B). To our knowledge, this work represents the first in-depth description of the etiology of ciliopathic micrognathia.

### NCC-derived skeletal precursors are highly sensitive to ciliopathic insults

Cell proliferation and ciliogenesis both require centriolar function and are thus inextricably linked. Previous studies solidified this relationship, reporting that knockdown of the ciliary protein IFT88 promoted cell cycle progression and proliferation (Robert et al., 2007). Our current study revealed that NCC-derived skeletal progenitors in the MNP were specifically affected, as seen by increased cell proliferation in the SOX9+ population of *ta*^*2*^ MNPs. Interestingly, our previous studies examining cell proliferation in early, undifferentiated NCC populations of the frontonasal prominence did not reveal any alteration in proliferation rates between control and *ta*^*2*^ embryos (Schock et al., 2015). Thus, these data beg the question: are NCC-derived skeletal progenitors more sensitive to a ciliopathic insult than other cell types? There is some precedent for the idea of cell specific sensitivities. Treacher-Collins syndrome (TCS) is a congenital craniofacial disorder characterized by hypoplasia of facial bones, high arched palate, and ear defects. Seminal studies addressing the etiology of TCS determined that NCCs, due to their high energetic demands as highly proliferative and migratory multipotent cells, were especially sensitive to deficient ribosome biogenesis and subsequent cellular stress (Dixon et al., 2006; Jones et al., 2008). Studies on various cell types are beginning to uncover a connection between the cilium and energy homeostasis (Arsov et al., 2006; Collin et al., 2005; Davenport et al., 2007; Marion et al., 2009; Pampliega et al., 2013; Tang et al., 2013). Additionally, the process of osteogenic differentiation, and maturation of osteoblasts in particular, has a heavy requirement of oxidative phosphorylation and glycolysis (Guntur et al., 2014; Regan et al., 2014). Thus, the lack of primary cilia on skeletal progenitors could result in a similar type of cellular stress, preventing *ta*^*2*^ NCCs from executing proliferation and differentiation programs that allow for proper skeletogenesis in the developing MNP.

### Determining the molecular etiology for ciliopathic micrognathia

We have previously shown that all facial prominences of *ta*^*2*^ mutants have perturbed GLI processing and subsequent disruptions in Hh signal transduction (Chang et al., 2014). The Hh pathway is a key player in skeletal differentiation, as mouse models with impaired expression of Hh ligands and their obligate transducer Smoothened possess a variety of craniofacial bone and cartilage defects (Abzhanov et al., 2007; Billmyre and Klingensmith, 2015; Jeong et al., 2004; Lenton et al., 2011; Xu et al., 2019). Furthermore, mice with impaired expression of GLI transcription factors also present with craniofacial skeletal defects (Elliott et al., 2020; Mo et al., 1997). GLIs are post-translationally processed into activator (GLIA) or repressor (GLIR) isoforms at the primary cilium (Goetz and Anderson, 2010; Haycraft et al., 2005; Liu et al., 2005) and loss of properly processed GLI isoforms results in altered transcription of Hh pathway target genes (Chang et al., 2014).

ChIP data investigating GLI2 and GLI3 binding sites in the developing MNP of mice demonstrate that these transcription factors have possible regulatory relationships with skeletogenic transcription factors. GLI peaks were observed near the transcriptional start site of *HAND2* and the middle of the *RUNX2* locus (Elliott et al., 2020). Further studies reported interactive roles for GLI proteins with skeletal transcription factors. GLI2 was reported to physically interact with RUNX2 to direct osteoblast differentiation (Shimoyama et al., 2007) and GLI3 was reported to utilize HAND2 as a co-factor to drive the mandibular patterning and osteogenic transcriptional programs (Elliott et al., 2020). These studies, together with the data we presented herein, suggest that impaired osteogenic differentiation in the *ta*^*2*^ MNP could be downstream of disrupted GLI-mediated Hh signaling. Future experimentation, including attempting to rescue skeletal defects with ectopic expression of processed GLI proteins, will need to be carried out to test this hypothesis.

Other signaling pathways, such as the Wnt, Fibroblast Growth Factor (FGF), and Bone Morphogenetic Protein (BMP) pathways, are also known to be involved in skeletal development (Jiang et al., 2014; Merrill et al., 2008; Mina et al., 2007). However, the mechanisms by which these pathways require cilia-dependent signaling transduction is less clear. The role of the cilium in canonical Wnt signaling has been heavily disputed. Initial studies demonstrated that numerous Wnt pathway components, such as Inversin, Vangl2 and Apc, localized to the ciliary axoneme or basal body, suggesting a role for primary cilia in Wnt signal transduction (Morgan et al., 2002; Ross et al., 2005; Simons et al., 2005). Subsequent studies demonstrated that the cilium functions to repress Wnt signaling as loss of cilia correlated with increased Wnt activity and Wnt target gene expression (Corbit et al., 2008; Lancaster et al., 2011). However, concurrent studies in mice and zebrafish that lack various ciliary proteins retained normal levels of canonical and non-canonical Wnt signaling transduction (Huang and Schier, 2009; Ocbina et al., 2009). Further, in the NCC-derived facial mesenchyme, conditional loss of the ciliary protein Kif3a in NCCs did not appear to alter Wnt activity (Brugmann et al., 2010). Recently, it was found that deletion of Kif3a plays a role in Wnt signaling in a ciliary-independent fashion by activating the pathway in an autocrine manner (Kim et al., 2016). If and how C2CD3 regulates Wnt activity in the *ta*^*2*^ during craniofacial skeletal development has not yet been described. Our future research will examine the impact loss of C2cd3-mediated ciliogenesis has on Wnt signaling.

BMP and FGF signaling pathways are also required for mandibular skeletogenesis (Ashe et al., 2012; Merrill et al., 2008; Mina et al., 2007), but understanding of how these signals are affected in ciliopathic backgrounds is poorly understood. The ciliopathic *Fuz*^*−/−*^ mouse mutant exhibits craniofacial features that closely resemble FGF hyperactivation syndromes, such as craniosynostosis and high arched palate (Tabler et al., 2013). Additionally, our previous studies with the *ta*^*2*^ mutant demonstrated that the developing MNP and maxillary prominence have increased FGF signaling (Schock et al., 2015). However, the mechanistic link between cilia and the FGF signaling pathway has not yet been established. Additionally, it is unclear if and how a BMP signal is transduced through the primary cilium (reviewed in (Kaku and Komatsu, 2017)). Our future goals include gaining a broader understanding of how other signaling pathways essential for skeletal development are affected in ciliary mutants.

### Pharmacological intervention could alleviate some aspects of ciliopathic micrognathia

Bone is constantly remodeled through bone deposition and bone resorption. Disturbances in the skeletal environment caused by excessive bone remodeling lead to decreased jaw length and density (Ealba et al., 2015; Feng and McDonald, 2011). As bone remodeling requires both bone deposition and bone resorption, defects in either program can lead to bone remodeling diseases such as osteoporosis and osteopetrosis (Feng and MacDonald, 2011). Additionally, it has been long established that osteoblasts and osteoclasts communicate to control bone remodeling through physical interaction and release of various growth factors and chemokines (Matsuo and Irie, 2008); however, how primary cilia mechanically or molecularly contribute to this communication has not yet been established. Previous studies have reported that primary cilia may play an antiresorptive role in osteocytes through increasing the OPG/RANKL ratio (Malone et al., 2007). While our studies confirm this finding as loss of cilia resulted in increased bone resorption in the *ta*^*2*^ mandible, future work on the molecular and mechanical control of bone resorption by primary cilia are necessary.

The discovery that bone remodeling contributes to ciliopathic micrognathia is a significant finding as it opens a potential avenue of therapeutic intervention. Studies have shown that bone resorption can be pharmacologically induced or depleted with recombinant forms of OPG and RANKL in avian models (Ealba et al., 2005). Additionally, bisphosphonates have previously been utilized to decrease bone resorption by preventing osteoclast formation and inducing osteoclast apoptosis (Boonekamp et al., 1986; Lowik et al., 1988; Sato and Grasser, 1990). Lastly, specific αNAC polypeptides have the potential to induce osteoblastic maturation *in vivo* (Meury et al., 2010). Treating ciliopathic micrognathia models, including the *ta^2^*, with these compounds during discrete temporal windows of remodeling is a focus of our ongoing research.

### Mechanosensory mechanisms may also contribute to ciliopathic micrognathia

It is well-established that increased mechanical loading stimulates bone formation (reviewed in (Robling and Turner, 2009)), and numerous studies have demonstrated a role for primary cilia as mechanosensory organelles during skeletal development (Leucht et al., 2013; Malone et al., 2007; Temiyasathit and Jacobs, 2010; Temiyasathit et al., 2012). Conditional loss of cilia in osteoblasts and osteocytes caused reduced bone formation due to reduced loading (Temiyasathit et al., 2012), decreased osteogenic response during fracture healing (Leucht et al., 2013), and inhibition of fluid flow-induced expression of osteogenic markers (Malone et al., 2007). When cilia on murine osteoblasts were removed with chloral hydrate, there was a lack of mineral deposition as a response to oscillatory fluid flow, demonstrating the importance of cilia for bone development (Delaine-Smith et al., 2014).

Several studies have suggested that a key role of the primary cilium is to function as chemical sensor for extracellular Ca^2+^ (Delling et al., 2013; Nauli et al., 2016; Zayzafoon, 2006). Primary cilia possess membrane-based ion channels including PKDL1/TRPP3, which allow for intracellular flux of Ca^2+^ (DeCaen et al., 2013; Nauli et al., 2016). Although this theory is controversial (Delling et al., 2016), this is an interesting direction for future experimental work given that bone mineralization requires Ca^2+^. Ca^2+^ is essential for bone mineralization as it precipitates with inorganic phosphate to form hydroxyapatite crystals in collagen-rich extracellular matrix (reviewed in (Murshed, 2018)). Low Ca^2+^ intake and incorporation leads to severe skeletogenic disorders, such as osteoporosis and osteopenia, that are presented with low osteoblast maturation and increased bone remodeling (Anderson, 1996; Dymling, 1964). Could C2cd3-mediated loss of cilia prevent Ca^2+^ processing or integration into skeletal elements? This question is particularly intriguing for the fact that bone deposition, differentiation and remodeling are connected to Ca^2+^ signaling (Delling et al., 2013; Nauli et al., 2016; Zayzafoon, 2006) and that C2cd3 contains several C2 Ca^2+^-dependent domains that are very poorly understood. Future studies will help to address if and how Ca^2+^ integration is affected in ciliopathic mutants and will be crucial for the understanding of how micrognathia and other ciliopathic skeletal disorders can be treated.

## Materials and Methods

### Avian embryo collection, genotyping and tissue preparation

Fertilized control and *ta*^*2*^ eggs were supplied from University of California Davis. Embryos were incubated at 38.8 °C for 5 – 13 days and then harvested for analysis. All embryos collected were staged according to the Hamburger-Hamilton staging system and genotyped as previously described (Chang et al., 2014; Hamburger and Hamilton, 1951). Embryos were fixed in 4% paraformaldehyde (PFA) overnight at 4°C, unless noted otherwise.

### Whole-mount skeletal staining

Whole-mount skeletal staining was performed as previously described (Rigueur and Lyons, 2014) with several modifications. Briefly, embryos were collected in PBS and fixed in 95% EtOH overnight. To remove excess adipose tissue, embryos were transferred to acetone overnight. To stain cartilage, embryos were placed in 0.03% Alcian Blue solution (Sigma-Aldrich A5268) for 2 hours. Following cartilage staining, samples were washed with 7 parts EtOH and 3 parts acetic acid until the solution was clear and incubated in 95% EtOH overnight. To stain bone, samples were incubated in 0.005% Alizarin Red S (Sigma-Aldrich A5533) in 1% KOH for 3 hours at room temperature and cleared in 1% KOH. Once cleared, samples were incubated in 50% glycerol 50% KOH solution. For imaging and long-term storage, samples were kept in 100% glycerol. Stained specimens were imaged using a Leica M165 FC stereo microscope system.

### Skeletal element analysis

Mandibular and ceratobranchial length measurements were performed using Leica LAS X software. Mandibles were measured from the distal tip of the dentary to the proximal edge of the surangular for each side of the mandible and then averaged. For individual skeletal element volumetric analysis, individual bones were traced in Fiji (Schindelin et al., 2012) and areas of each bone were measured and calculated. Student’s t-test was used for statistical analysis. *P*<0.05 was determined to be significant.

### Micro-computed tomography (Micro-CT)

HH36 and HH39 mandibles were harvested, fixed in 4% PFA, and stained with Alizarin red for further processing at the Preclinical Imaging Core in the University of Cincinnati Vontz Center for Molecular Studies. DICOM files were processed, with voxel quantification of the surface and volume areas performed in iMaris 9.3 (Bitplane – Oxford Instruments).

### Whole-mount immunofluorescence

Whole mandible immunofluorescence was carried out as previously described with minor modifications (Williams et al., 2018). Triton-X concentration was increased to 1% and embryos were incubated in blocking solution supplemented with 5% normal goat serum. Cleaved Caspase-3 (1:100, Cell Signaling 9661) and Phospho-Histone H3 (1:200, Cell Signaling 9706) primary antibodies were used. The secondary antibodies used were Goat anti-Mouse-568 (1:250, ThermoFisher A-11004), Goat anti-Rabbit-647 (1:250, ThermoFisherA21244), and Sox9-Conjugate AlexaFluor-488 (1:100 - Cell Signaling – 94794). Nuclei counterstaining was carried out by incubation in 1 μg/ml DAPI in PBS + Triton 1% for 2 hours and processed for clearing with Ce3D solution overnight (Li et al., 2017).

### RNAScope *in situ* hybridization

HH29 heads and HH39 mandibles were fixed in 4% PFA at 4°C for 16-24 hours. HH39 mandibles were decalcified in 19% EDTA solution at 4°C for 5 days. Samples were dehydrated in an ethanol series, washed in xylene, embedded in paraffin, and sectioned at 8μm thickness. Transcripts of *RUNX2* (ACD 571591), *HAND2* (ACD 571571-C2), and *DLX5* (ACD 571561) were detected using the RNAscope Multiplex Fluorescent V2 kit per manufacturer’s instructions. Transcripts of *SPP1* (ACD 571601), *ALPL* (ACD 837811), and *MMP13* (ACD 571581) were detected using RNAscope 2.5 HD Duplex Assay (ACD) per manufacturer’s instructions. Briefly, slides were baked in a hybridization oven for 1 hr at 60°C, deparaffinized using xylene, dehydrated using 100% EtOH, and allowed to dry completely at room temperature. Endogenous peroxidase activity was quenched using provided hydrogen peroxide and washed using distilled water. Target retrieval was performed using provided Target Retrieval Buffer in an Oster steamer for 15 min, washed in distilled water, dehydrated in 100% EtOH, and dried at 60°C. Slides were treated with ACD Protease Plus in an HybEZ II Oven for 30 min at 40°C. Probes were hybridized for 2 hr at 40°C in an HybEZ II Oven. Amplification steps were performed as described by manufacturer. Signal development for *RUNX2*, *HAND2*, and *DLX5* were carried out using Cyanine 3 (PerkinElmer NEL752001KT) diluted 1:500 in RNAscope Multiplex TSA Buffer. Signal amplification for *SPP1*, *ALPL*, and *MMP13* were carried out using provided HRP-based Green chromogen. Slides were counterstained and mounted per manufacturer’s instructions and imaged using a Leica DM5000B upright microscope system.

### Fluorescent *in situ* hybridization

Fluorescent whole mount *in situ* hybridization (FISH) was performed using digoxigenin-labeled riboprobes as previously described (Denkers et al., 2004). Antisense riboprobes against *DLX5* (NM_204159.1, primers used F-5’-ATCAGGTCCTCCGACTTCCA-3’ and R-5’-ATACGACTCACTATAGGGGATTTTCACCTGCGTCTGCG – 3’), *HAND2* (NM_204966.2 – primers used F-5’ – CGAGGAGAACCCCTACTTCC – 3’ and R-5’ – TAATACGACTCACTATAGGGCCTGTCCGCCCTTTGGTTT – 3’) and *RUNX2* (NM_204128.1 – primers used F-5’–CGCATTCCTCATCCCAGTAT–3’ and R-5’-TAATACGACTCACTATAGGGTATGGAGTGCTGCTGGTCTG -3’) were designed to range from 600 to 800bp. TSA Plus HRP-Fluorescence kits were used for both Cy3 and Cy5 fluorescence channels (PerkinElmer NEL752001KT). For simultaneous SOX9 immunofluorescence, embryos were washed with TNT and incubated overnight in 5% normal goat serum in TNT. Nuclei were counterstained with DAPI.

### Quantitative reverse transcriptase PCR

RNA was extracted using TRIzol reagent (Invitrogen) and cDNA was synthesized using SuperScript III (Invitrogen). HH39 mandibles were first frozen with liquid nitrogen and ground using a mortar and pestle to ensure homogenous extraction from bone. SYBR Green Supermix (Bio-Rad) and a Quant6 Applied Biosytems qPCR machine were used to perform quantitative real-time PCR. All the genes were normalized to GAPDH expression. Student’s *t*-test was used for statistical analysis. *P*<0.05 was determined to be significant.

## Supplemental Information

Supplemental Information includes three Figures.

## Author Contributions

C.L.B.P. performed and analyzed qRT-PCR quantification, immunofluorescence, fluorescent *in situ* hybridization, and microCT analysis. E.C.B. performed Alizarin red staining of the mandible, mandible length and volume measurements, and RNAscope *in situ* hybridization and analyzed data together with the other authors. M.A.P. performed the Alizarin red/Alcian blue staining of the mandibular and hyoid elements and initial characterization of the mandibular phenotype. S.A.B. acquired funds, idealized the project, discussed data and interpreted results, and wrote the manuscript with input from C.L.B.P. and E.C.B.

## Acknowledgements

We would like to thank Mary Delany and the UC Davis Avian Facility, Jackie Pisenti and Kevin Bellido for maintenance and husbandry of the *talpid*^*2*^ colony. Technical assistance was given by Dr. Matt Kofron and Evan Meyer for image acquisition and analysis (Confocal Imaging Core – CCHMC) and Dr. Lisa Lemen for microCT acquisition and analysis (Preclinical Imaging Core – University of Cincinnati). We would also like to thank members of the Brugmann lab for helpful comments and feedback. This study was funded by the National Institutes of Health (R35 DE027557) and Shriners Hospital for Children (543938) to S.A.B.

---

**Supplementary Fig. 1.**
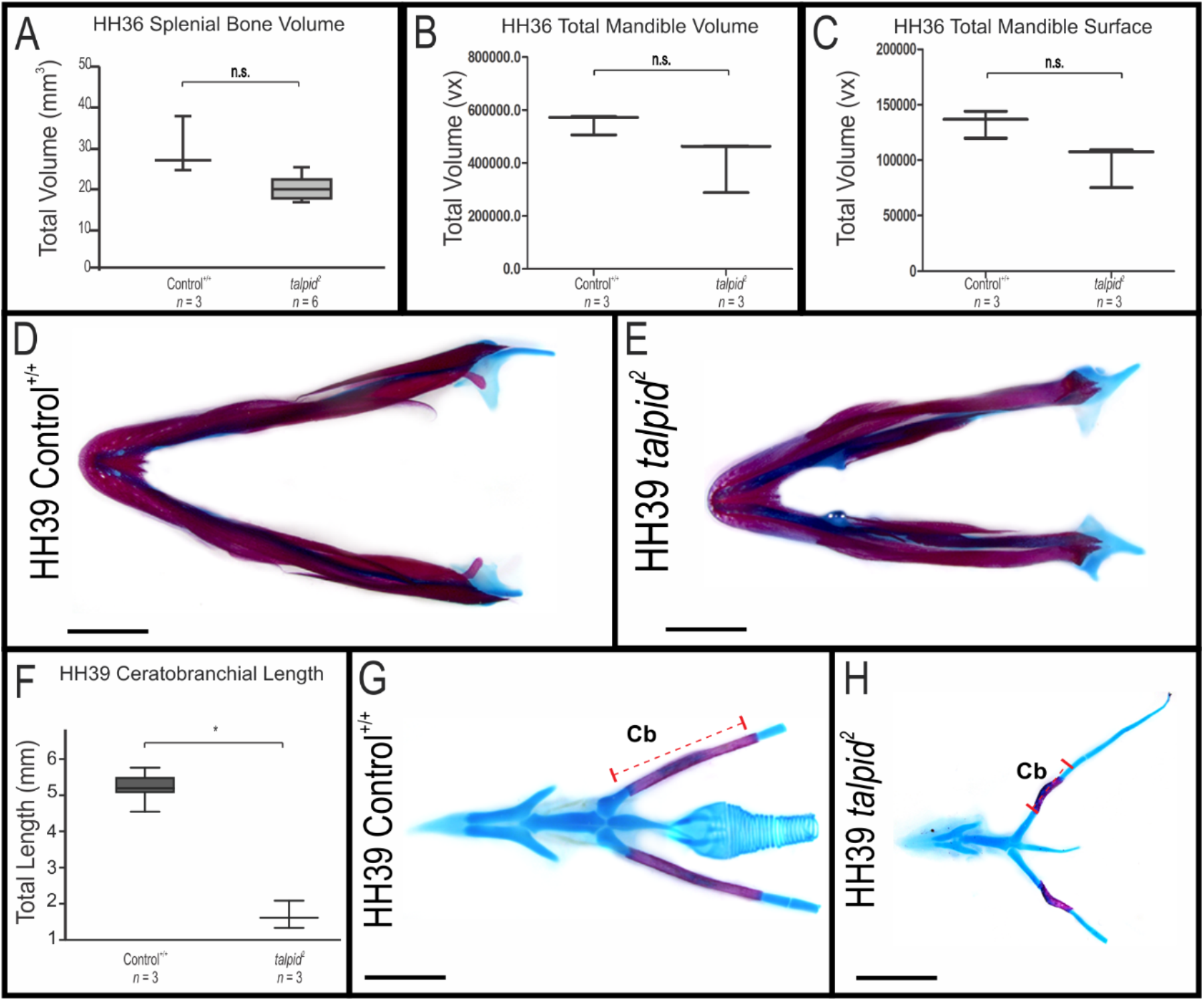
Measurements for *ta*^*2*^ mandibles. (A) Volume measurements of HH36 control^+/+^ and *ta*^*2*^ splenial bone. (B-C) HH36 total mandible (B) volume and (C) surface area of control^+/+^ and *ta*^*2*^ mandibles. (D-E) Alcian blue and Alizarin red staining of HH39 control^+/+^ and *ta*^*2*^ mandibular skeletons (n = 7). (F) Length measurements of HH39 control^+/+^ and *ta^2^* ceratobranchial bones. (G-H) Alcian blue and Alizarin red staining of HH39 control^+/+^ and *ta*^*2*^ hyoid skeletons. Cb: Ceratobranchial, Vx: Voxels. Data are mean ± s.d. Scale bars are 1cm (D-E, G-H). Statistical analysis was performed by student’s t-test (* denotes *P*< 0.001).

**Supplementary Fig. 2.**
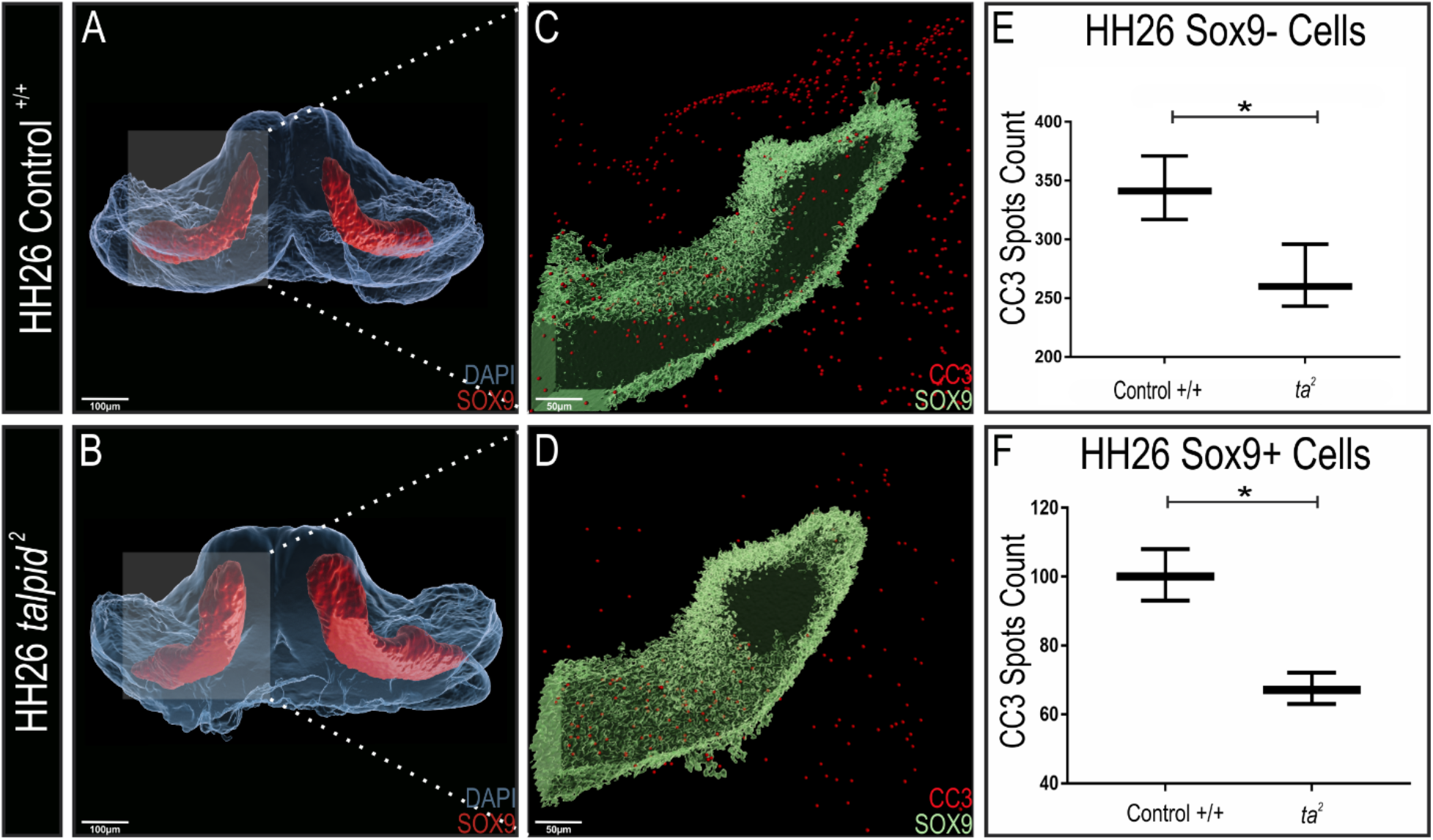
Cell death is decreased in *ta*^*2*^ MNPs. (A-B) Wholemount MNP staining for SOX9 (red) and DAPI (blue) in HH26 control^+/+^ and *ta*^*2*^ MNPs. (C-D) Puncta projection for CC3 (red) and SOX9 (green) in HH26 (C) control^+/+^ and (D) *ta*^*2*^ MNPs. (E) CC3 puncta count in the SOX9-cell population of control^+/+^ and *ta*^*2*^ MNPs (n = 3). (F) CC3 puncta count in the SOX9+ cell population of HH26 control^+/+^ and *ta*^*2*^ MNPs (n = 3). Data are mean ± s.d. Statistical analysis was performed by student’s t-test (* denotes *P*< 0.005).

**Supplementary Fig. 3.**
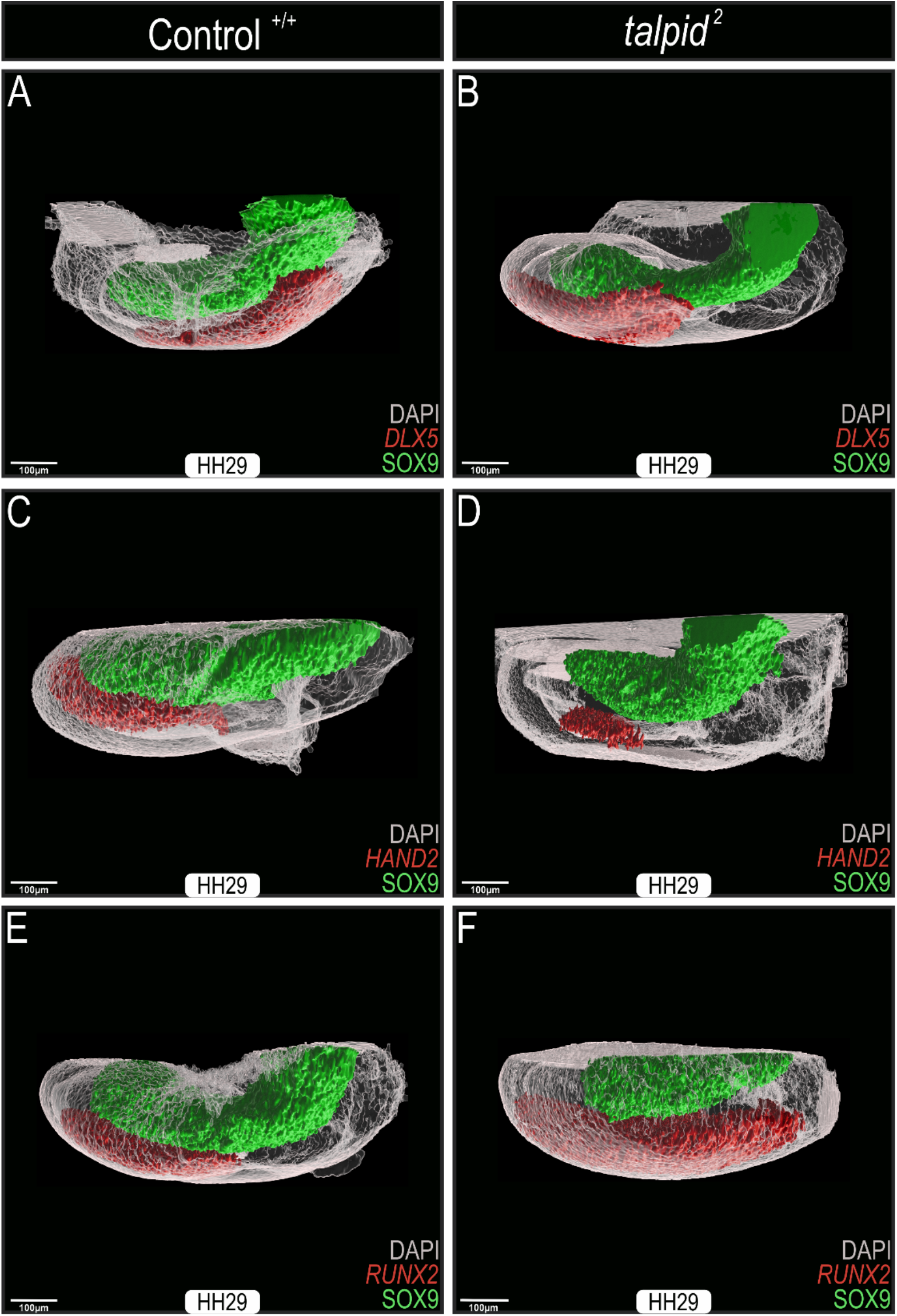
Orthogonal projections of HH29 control^+/+^ and *ta*^*2*^ MNPs. (A-B) Lateral view of *DLX5* and SOX9 stained control^+/+^ and *ta*^*2*^ MNPs at HH29. (C-D) Lateral view of *HAND2* and SOX9 stained control^+/+^ and *ta*^*2*^ MNPs at HH29. (E-F) Lateral view of *RUNX2* and SOX9 stained control^+/+^ and *ta*^*2*^ MNPs at HH29. Z-stacks were taken at every 5μm, ranging from 600 to 800μm.

